# DAnIEL: A User-Friendly Web Server for Fungal ITS Amplicon Sequencing Data

**DOI:** 10.1101/2021.04.12.437814

**Authors:** Daniel Loos, Lu Zhang, Christine Beemelmanns, Oliver Kurzai, Gianni Panagiotou

## Abstract

Trillions of microbes representing all kingdoms of life are resident in, and on, humans holding essential roles for host development and physiology. The last decade over a dozen online tools and servers, accessible via public domain, have been developed for the analysis of bacterial sequences, however, the analysis of fungi is still in its infancy. Here we present a web server dedicated to the comprehensive analysis of the human mycobiome for (i) translating raw sequencing reads to data tables and high-standard figures; (ii) integrating statistical analysis and machine learning with a manually curated relational database; (iii) comparing the user’s uploaded datasets with publicly available from the Sequence Read Archive. Using 2,048 publicly available ITS samples, we demonstrated the utility of DAnIEL web server on large scale datasets and show the differences in fungal communities between human gut, skin, nasopharynx, and oral body sites.

## Introduction

Metagenomics provides a comprehensive view about microbial community structure. Previous studies have revealed many insights about the diversity, composition, and interaction patterns of bacterial communities. Fungi are a neglected but very important kingdom, due to the important role they play in many human diseases [1] The number of publications in PubMed related to the mycobiome is exponentially growing and increased more than 17-fold in the past five years. Fungal metagenomics is becoming essential part for comprehensive human host studies and should be accessible to the whole scientific community without the need of laborious and time-consuming efforts. We present here DAnIEL (Describing, Analyzing and Integrating fungal Ecology to effectively study the systems of Life) the only web server available that covers the whole workflow of ITS analysis beginning from raw reads to publication ready figures and tables, contains a relational database for biological evaluation of statistical findings and allows comparative analysis with public available mycobiome datasets. For all steps, a summary of methods and results including citations is provided and interactive plots can be created tailor-made. DAnIEL is freely available at https://sbi.hki-jena.de/daniel.

## Design and Implementation

### Overview

The workflow is illustrated in Figure 1. Raw reads can be uploaded in compressed FASTQ format. Optionally, read runs from the NCBI Sequence Read Archive (SRA) can be added by either selecting from the 700 existing cohorts of the DAnIEL database or by entering their accessions directly. Metadata about the samples can be uploaded in CSV or Excel file format, if statistical analysis is needed. Parameter sets for tweaking the workflow can be created e.g. to filter features by abundance or to trim custom primer sequences. A comprehensive documentation about parameters to tweak the workflow and a tutorial is available on the DAnIEL web server. To annotate significant features, we constructed a manually curated database containing 1,669 fungal interactions with diseases, bacteria species and immune components retrieved from 761 published papers. This database is used by the web server for annotation of significant differently abundant or correlated taxa. Furthermore, we incorporated a list of clinical samples of species involved in a suspicious fungal infection from the German National Reference Center for Invasive Fungal Infections (NRZMyk).

**Fig 1.**
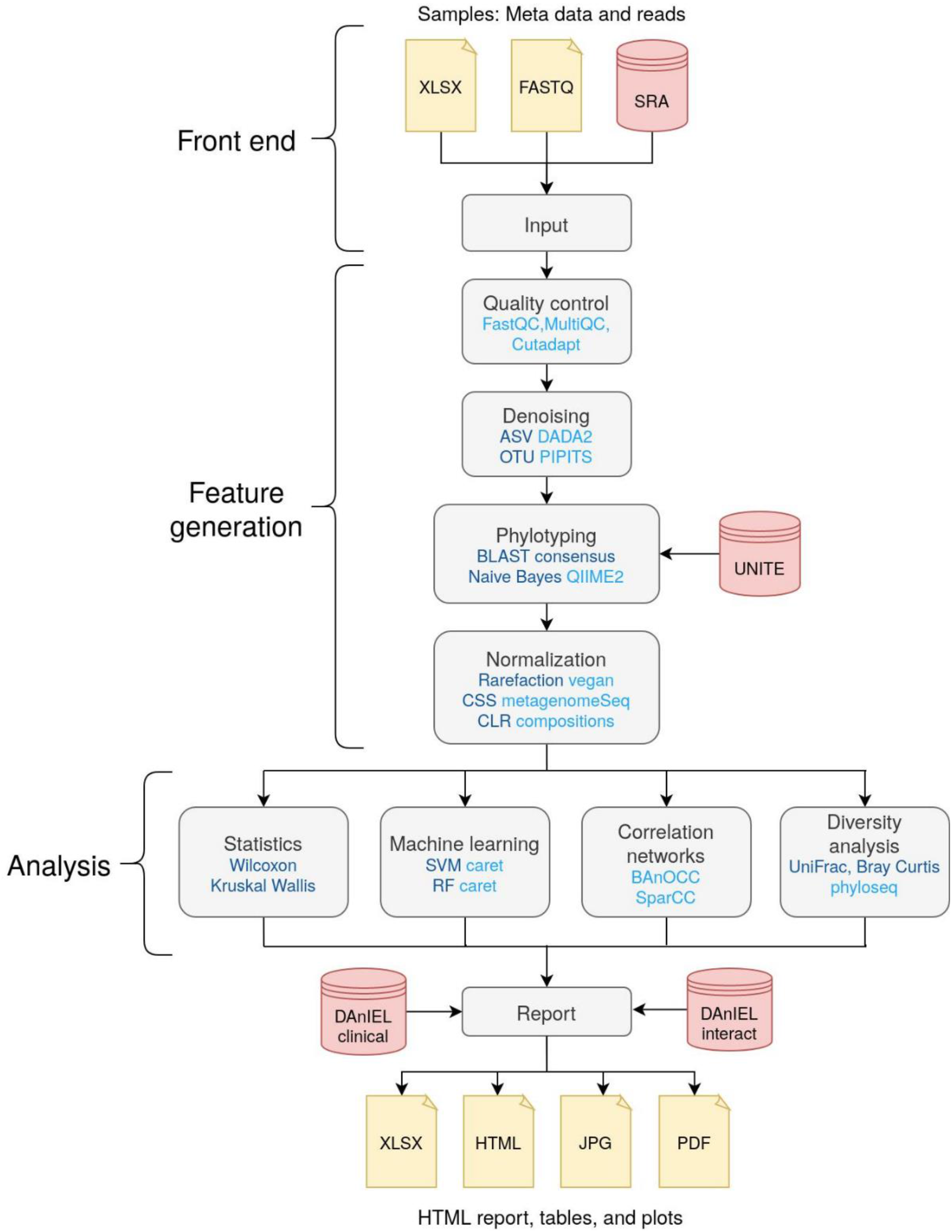
Overview about the workflow of DAnIEL web server. Methods and tools are shown in dark and light blue, respectively. Databases are shown in red. Feature generation and analysis are part of the back end. Taxa are augmented using our relational database of clinical samples (DanIEL clinical) and interactions reported in the literature (DanIEL interact).

### Feature generation

Features are generated from the raw reads provided. Samples are demultiplexed if necessary, according to the barcode mapping provided in the metadata table. External samples are downloaded from the NCBI Sequence Read Archive using grabseqs [2]. Quality control (QC) is performed afterwards. FastQC and MultiQC are used to monitor sequencing errors [3]. Cutadapt is used to trim primer and adapter sequences [4]. Samples can be excluded from downstream analysis using various criteria such as minimum number of quality-controlled reads or base quality tests specified by FastQC. Representative biological sequences are created from quality-controlled reads via denoising. Either OTUs or amplicon sequence variants (ASVs) can be called using PIPITS [5] or DADA2 [6], respectively. Taxonomy of denoised sequences is assigned using either Naive Bayes or BLAST consensus approach of QIIME2 [7]. Abundance counts are pooled at any given taxonomic rank and filtered by abundance and prevalence. Lastly, pooled counts are normalized using methods aware of different library sizes like rarefaction or cumulative sum scaling (CSS), as implemented in the R packages vegan and metagenomeSeq, respectively [8,9]. Centered log-ratio (CLR) normalization is used by default to account for the compositionality. The generated features are used in downstream analysis to infer biological insights.

### Benchmarking parameters

The performance of taxonomic profiling was evaluated using simulated reads. Grinder was used to simulate 10 samples in which each of them consists of 500k paired-end (PE) reads [22]. Primers ITS1 and ITS2 targeting ITS subregion were utilized to get 100 different sequences following an exponential abundance distribution. The same database was used for simulation and training the classifiers to enable a fair comparison (UNITE 8.2 dynamic) [23]. To simulate biological variability, a uniform mutation rate of 1% was incorporated in which substitutions were four times more likely than insertions or deletions. DAnIEL was run on the data with different methods for denoising (DADA2 and PIPITS) and taxonomic classification (BLAST consensus and Naive Bayes). Benchmarking performance of profiling abundance was based on [24] using counts pooled at genus rank. Briefly, the L2 norm between the measured and the true abundance profile was calculated for each sample. Furthermore, differences of abundances were calculated for each sample and taxon separately. Pure taxon occurrence was evaluated by counting samples in which a taxon was both measured and simulated. Precision, Sensitivity, Specificity and F1 score were calculated based on this contingency table. Taxon occurrences were very specifically but less sensitively profiled (98% and 33% on average, respectively, see Figure S1). DADA2 outperformed PIPITS in all metrics. Naive Bayes classification outperformed the BLAST consensus approach in terms of specificity and precision but not in sensitivity. Most accurate abundance profiles were generated using DADA2 (See Figure S2). PIPITS underestimated many abundances which lead to increased distances in some samples especially in combination with the Naive Bayes classifier. The values were very similar for different phyla. Therefore, taxonomy seemed to have only little influence on the classification performance.

### Relational database generation

DAnIEL was initially run on three cohorts to retrieve a list of fungal species relevant for analyses when studying human samples: fecal samples from a publicly available dataset of human skin swab samples (N=203, ITS1, PRJNA286273 [10]) and *in-house* mycobiome datasets from stool samples of cancer patients (N=71, ITS2) and a healthy volunteers during an antibiotics intervention study (N=59, ITS2) were used. For each species found, we constructed a NCBI Entrez query to search for PubMed abstracts. Terms ‘disease’, ‘cytokine’, ‘immune system’, and ‘prokaryote’ and a limit of 20 papers per species were used to narrow down the focus of our subsequent manual curation. In total 1337 abstracts from these papers were reviewed to create a manually curated database of fungal interactions. Medical Subject Headings (MeSH) were used for annotations whenever applicable.

### Feature analysis

Diversity is calculated using the R packages vegan [8] and phyloseq [11]. Various methods including principal coordinates analysis (PCoA) and non-metric multidimensional scaling (NMDS) can be used to generate ordination plots. FastSpar implementation of the SparCC algorithm can be used to create correlation networks of co-abundant taxa [12,13]. Alternatively, BAnOCC can be chosen to account for the compositionality of NGS abundance data [14]. The correlation analysis can be executed for each sample group individually e.g. to compare networks of “case” and “control” samples. If a metadata table is provided, group-wise statistics are performed using Mann–Whitney U test for binary response variables and Kruskal–Wallis one-way analysis of variance in combination with Dunn’s post hoc test [15] for other nominal responses. Spearman’s rank correlation is used for continuous responses instead. Features significant in any of these tests are annotated with our manually curated database of fungal interactions and clinical samples. Machine learning is applied to categorical response variables using the R package caret [16]. Both random forest (RF) and support vector machines (SVMs) are used in combination with ANOVA filter and recursive feature selection. Best performing models according to the area under the receiver operating curve (AUC) in 5-fold cross-validation and feature importance scores are reported.

### Technical design

The overall pipeline of the DAnIEL web server consists of two parts: a front-end the user is interacting with to upload and visualize the data and a back-end workflow responsible for processing the uploaded data. The front-end of DAnIEL web server is implemented as a R shiny app. For visualization ggplot2 is used [17] and Rmarkdown to create summary reports. The back-end is built as a Snakemake workflow [18]. This allows to run the workflow separately on any Linux system including computing clusters. Conda and Docker are used to create reproducible environments for installing and running scripts and individual tools. A unique identifier will be assigned to each project to access the results later on. This also acts as a token for authentication. The tutorial consisting of 38 samples usually takes approximately half an hour wall time using 10 threads to be fully processed. Reports and visualizations can be accessed at the front-end once the corresponding step in the workflow has finished. This includes interactive plots and a summary consisting of findings, annotations, methods, and references in a single HTML file.

## Results

### Comparison to related software

A summary of the functionality of alternative tools for mycobiome analysis is shown in Table 1. QIIME2 is command-line focused, therefore it is not ideal for researchers without programming skills [7]. ITScan covers profiling of operational taxonomic units (OTUs), however it starts from quality-controlled sequences [19]. CloVR-ITS was designed for pyrosequencing data [20]. To the best of our knowledge, DAnIEL is the only web server available covering the whole workflow of ITS analysis beginning from raw reads to publication ready figures and tables, as well as, integration with a relational database for biological evaluation of statistical findings and comparative analysis with public available mycobiome datasets.

**Tab 1.** Functionality of software for ITS analysis. Features of DAnIEL among the similar tools QIIME2, mothur, CloVR-ITS, and ITScan are compared.

| Feature | DAnIEL | QIIME2 | mothur | CloVR-ITS | ITScan |
| --- | --- | --- | --- | --- | --- |
| Existing cohorts for comparative analysis | + | - | - | - | - |
| Quality control | + | + | + | + | - |
| OTU/ASV profiling | +/+ | +/+ | +/- | +/- | +/- |
| SparCC/BAnOCC correlation | +/+ | +/- | +/- | -/- | -/- |
| Relational database | + | - | - | - | - |
| GUI | + | - | + | + | + |
| Webserver | + | - | - | - | + |

### Case study

To demonstrate the functionality of the DAnIEL web server, we did a meta-analysis of publicly available human mycobiomes. NCBI SRA was queried for Illumina human ITS amplicon runs. This yielded 2,048 samples that underwent a sanity check by aligning the first 1000 reads to the UNITE database using BLASTN (min. identity of 98.5%, max. e value of 10^−50^) [21]. Samples having less than 1% of the sampled reads aligned or lacking body site information were discarded. Body sites containing less than 50 samples were also not considered. Adapters and common primers as listed by UNITE were removed using cutadapt [4]. Feature counts were pooled at genus level with a minimum prevalence of 5% of samples in any body site group. DAnIEL workflow was run for ITS1 and ITS2 samples separately with default parameters. In total 457 ITS1 and 663 ITS2 samples passed feature generation and were available for downstream analysis. This included 414 fecal, 182 oral, 266 nasopharynx, and 258 skin samples. Alpha diversity of skin samples at the level of denoised ASVs was significantly lower compared to those taken at oral sites or the nasopharynx (Figure 2, Kruskal-Wallis *p* < 2.2 * 10^−16^, Dunn’s test *p* < 10^−16^ for all post hoc tests) using both Chao1 and Shannon indices. The overall mycobiome structure at genus level differed significantly in both ITS regions by body site (Figure 2, Bray-Curtis dissimilarity, Adonis PERMANOVA, *p* < 0.001). Within group beta diversity, using Bray-Curtis dissimilarity, was significant for all body sites (Kruskal-Wallis and Dunn’s test, *p* < 0.001) expect the comparison of mouth to gut samples. Network topology of co-abundant genera correlations differed significantly in all comparisons (Figure 2, Node degree and betweenness centrality, SparCC, Kruskal-Wallis, and Dunn’s test). SVM and RF were used to build classifiers able to distinguish between two body sites using CLR normalized abundances at genus level. Recursive feature elimination improved all classifiers in the ITS2 dataset. Best performing models according to the AUC in 5-fold Cross-Validation were selected. Average AUC ranged from 90% discriminating between oral and skin samples to 98% discriminating oral and nasopharynx samples. Gini index was used to calculate feature importance for all final RF models. Subsequently DAnIEL relational database of manually curated interactions and NRZMyk infections were queried for species in any of the 10 most important genera. We found species belonging to these genera to be involved in 149 clinical samples of fungal infections from different organs and retrieved 805 interactions in which 314 are related to a disease, 54 to the immune system, 53 to a cytokine, and 50 to a body site. This indicates the importance of such databases in interpreting ML models.

**Fig 2.**
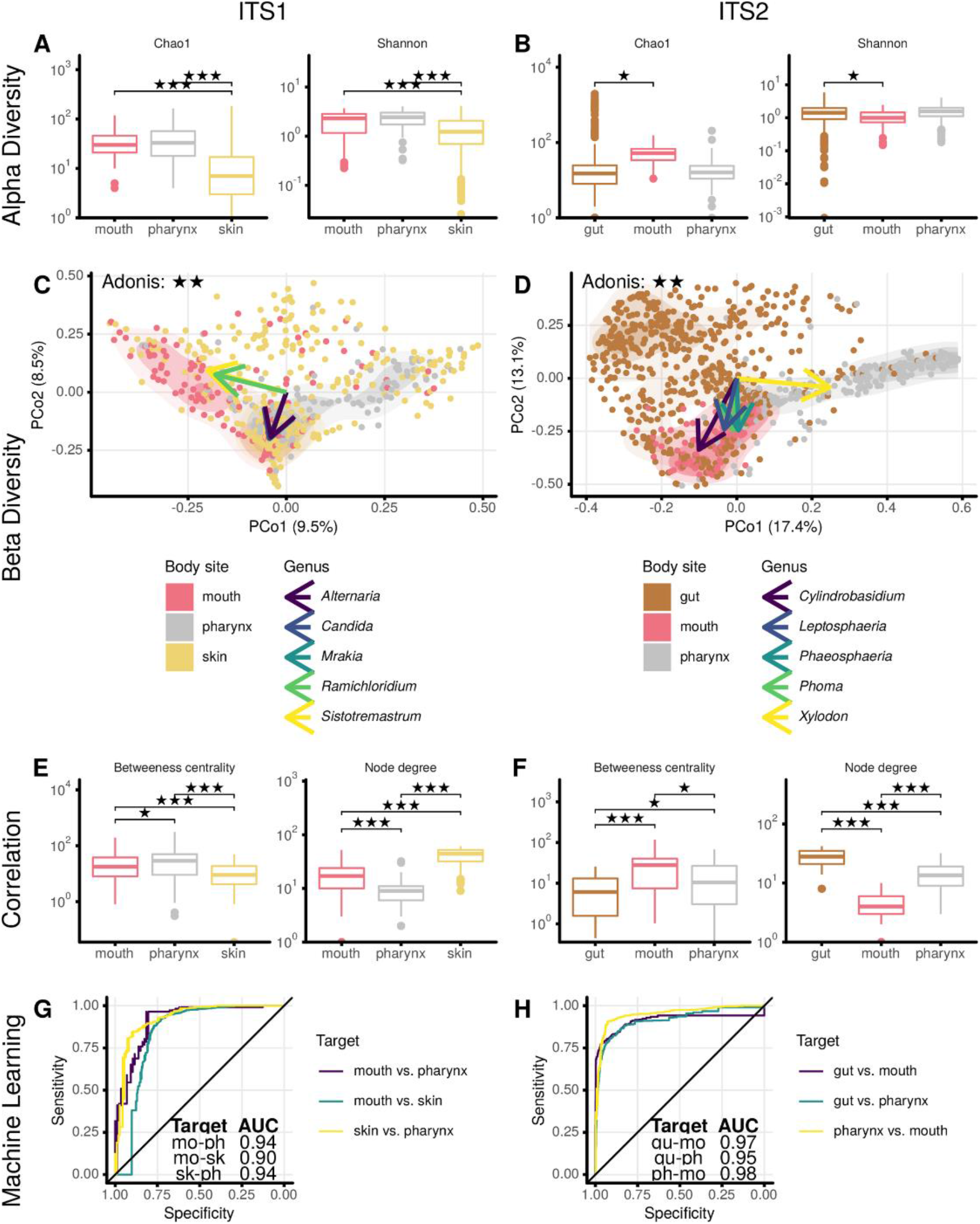
Results of the case study. Human ITS1 and ITS2 samples were processed with DAnIEL separately. Alpha Diversity (A and B), beta diversity (C and D), network topology of correlation networks (E and F), and performance of machine learning models (G and H) is shown. Mycobiomes differ significantly between different body sites (Bray-Curtis dissimilarity, Adonis PERMANOVA, *p* < 0.001)

## Availability and Future Directions

DAnIEL is freely available as a web service at https://sbi.hki-jena.de/daniel. There is no registration required. Instead, an ID token will be assigned to each project. Results will be available for 30 days. The source code is hosted at https://github.com/bioinformatics-leibniz-hki/DAnIEL, together with several tests and an example on how to use it, and is distributed under the BSD 2-Clause. There are also Docker images available to host own instances. All databases including reference sequences, existing cohorts and fungal interactions can be downloaded at https://doi.org/10.5281/zenodo.4073125. Using the Snakemake workflow engine, DAnIEL can be easily extended by other steps [18]. For instance, Picrust2 can be integrated to predict fungal function profiles [25]. Fungal taxonomic profiling profits from sequencing larger amplicons. Therefore, tools specifically designed for long read sequencing data of the third generation can be used for Quality Control to improve classification performance. Furthermore, updating the manually curated database and augmenting it with text mining approaches will improve the biological interpretation of significant taxa.

## Author contributions

The study was conceived by D.L. and G.P. The web server was designed by D.L and G.P and implemented by D.L. The relational database of fungal interactions was curated by L.Z, C.B and NRZMyk by O.K Existing projects were processed by D.L. The manuscript was written by D.L. and G.P. All authors approved the manuscript.

## Acknowledgments

We thank the members of NRZMyk for providing us the data about clinical samples with fungal infections.

## References

[1] Mukherjee JAR Pranab K. AND Chandra. Oral mycobiome analysis of hiv-infected patients: Identification of pichia as an antagonist of opportunistic fungi. PLOS Pathogens. 2014;10:1–17.

[2] Taylor LJ, Abbas A, Bushman FD. grabseqs: Simple downloading of reads and metadata from multiple next-generation sequencing data repositories. Bioinformatics. 2020.

[3] Ewels P, Magnusson M, Lundin S, Käller M. MultiQC: summarize analysis results for multiple tools and samples in a single report. Bioinformatics. 2016;32:3047–8.

[4] Martin M. Cutadapt removes adapter sequences from high-throughput sequencing reads. EMBnetjournal. 2011;17:10–12.

[5] Gweon HS, Oliver A, Taylor J, Booth T, Gibbs M, Read DS, et al. PIPITS: An automated pipeline for analyses of fungal internal transcribed spacer sequences from the illumina sequencing platform. Methods in ecology and evolution. 2015;6:973–980.

[6] Callahan BJ, McMurdie PJ, Rosen MJ, Han AW, Johnson AJA, Holmes SP. DADA2: High-resolution sample inference from illumina amplicon data. Nature methods. 2016;13:581–3.

[7] Bolyen E, Rideout JR, Dillon MR, Bokulich NA, Abnet CC, Al-Ghalith GA, et al. Reproducible, interactive, scalable and extensible microbiome data science using QIIME 2. Nat Biotechnol. 2019;37:852–857.

[8] Dixon P. VEGAN, a package of r functions for community ecology. Journal of Vegetation Science. 2003;14:927–930.

[9] Paulson JN, Stine OC, Bravo HC, Pop M. Differential abundance analysis for microbial marker-gene surveys. Nat Methods. 2013;10:1200–2.

[10] Leung MHY, Chan KCK, Lee PKH. Skin fungal community and its correlation with bacterial community of urban chinese individuals. Microbiome. 2016;4:46.

[11] McMurdie PJ, Holmes S. phyloseq: an R package for reproducible interactive analysis and graphics of microbiome census data. PLoS ONE. 2013;8:e61217.

[12] Watts SC, Ritchie SC, Inouye M, Holt KE. FastSpar: Rapid and scalable correlation estimation for compositional data. Bioinformatics (Oxford, England). 2019;35:1064–1066.

[13] Friedman J, Alm EJ. Inferring correlation networks from genomic survey data. PLoS computational biology. 2012;8:e1002687.

[14] Schwager E, Mallick H, Ventz S, Huttenhower C. A bayesian method for detecting pairwise associations in compositional data. PLoS computational biology. 2017;13:e1005852.

[15] Dunn OJ. Multiple comparisons using rank sums. Technometrics. 1964;6:241–252.

[16] Kuhn M. Building predictive models in r using the caret package. Journal of Statistical Software, Articles. 2008;28:1–26.

[17] Wickham H, Averick M, Bryan J, Chang W, McGowan L, François R, et al. Welcome to the tidyverse. Journal of Open Source Software. 2019;4:1686.

[18] Koster J, Rahmann S. Snakemake–a scalable bioinformatics workflow engine. Bioinformatics (Oxford, England). 2012;28:2520–2.

[19] Ferro M, Antonio EA, Souza W, Bacci M. ITScan: a web-based analysis tool for Internal Transcribed Spacer (ITS) sequences. BMC Res Notes. 2014;7:857.

[20] White JR, Maddox C, White O, Angiuoli SV, Fricke WF. CloVR-ITS: Automated internal transcribed spacer amplicon sequence analysis pipeline for the characterization of fungal microbiota. Microbiome. 2013;1:6.

[21] Camacho C, Coulouris G, Avagyan V, Ma N, Papadopoulos J, Bealer K, et al. BLAST+: architecture and applications. BMC Bioinformatics. 2009;10:421.

[22] Angly FE, Willner D, Rohwer F, Hugenholtz P, Tyson GW. Grinder: a versatile amplicon and shotgun sequence simulator. Nucleic Acids Res. 2012;40:e94.

[23] Nilsson RH, Larsson K-H, Taylor AFS, Bengtsson-Palme J, Jeppesen TS, Schigel D, et al. The unite database for molecular identification of fungi: Handling dark taxa and parallel taxonomic classifications. Nucleic acids research. 2019;47:D259–D264.

[24] Ye SH, Siddle KJ, Park DJ, Sabeti PC. Benchmarking Metagenomics Tools for Taxonomic Classification. Cell. 2019;178:779–794.

[25] Douglas GM, Maffei VJ, Zaneveld JR, Yurgel SN, Brown JR, Taylor CM, et al. PICRUSt2 for prediction of metagenome functions. Nat Biotechnol. 2020;38:685–688.

