## Supplementary Information for "DAnIEL: A User-Friendly Web Server for Fungal ITS Amplicon Sequencing Data"

[1]Systems Biology and Bioinformatics, Leibniz Institute for Natural Product Research and Infection Biology, Germany, [2]Chemical Biology of Microbe-Host Interactions, Leibniz Institute for Natural Product Research and Infection Biology, Germany, [3]Institute for Hygiene and Microbiology, University of Würzburg, Germany, [4]Systems Biology and Bioinformatics, School of Biological Sciences, Faculty of Science, The University of Hong Kong, China

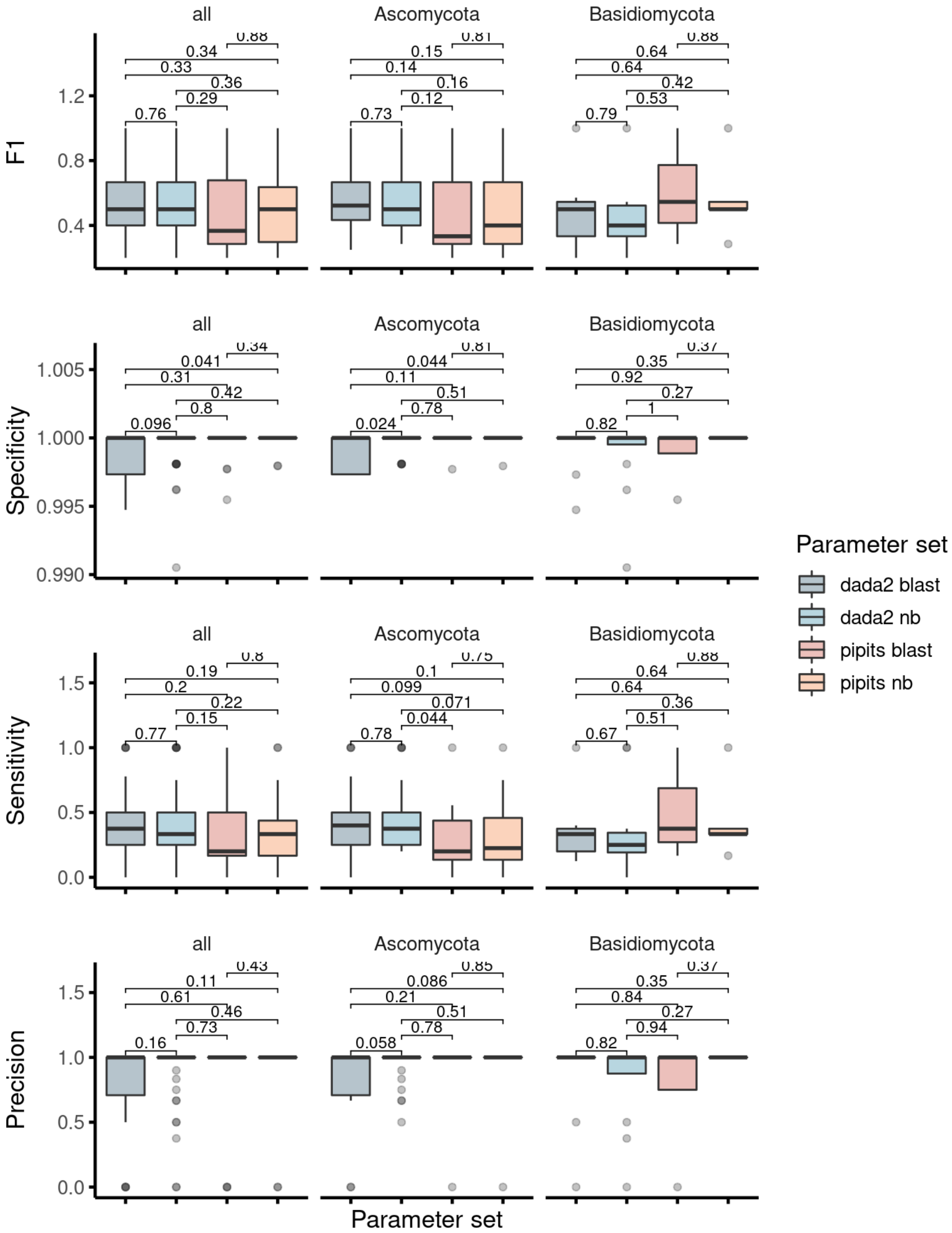


**Fig. S1:** Benchmarking taxon existence. DAnIEL was run on 10 samples consisting simulated reads using denoising methods DADA2 and PIPITS and classification methods BLAST consensus (blast) and Naïve Bayes (nb). Contingency tables were calculated by counting samples in which a taxon was both measured and simulated. DADA2 outperformed PIPITS in all metrics. NB classification outperformed BLAST in terms of specificity and precision but not in sensitivity.

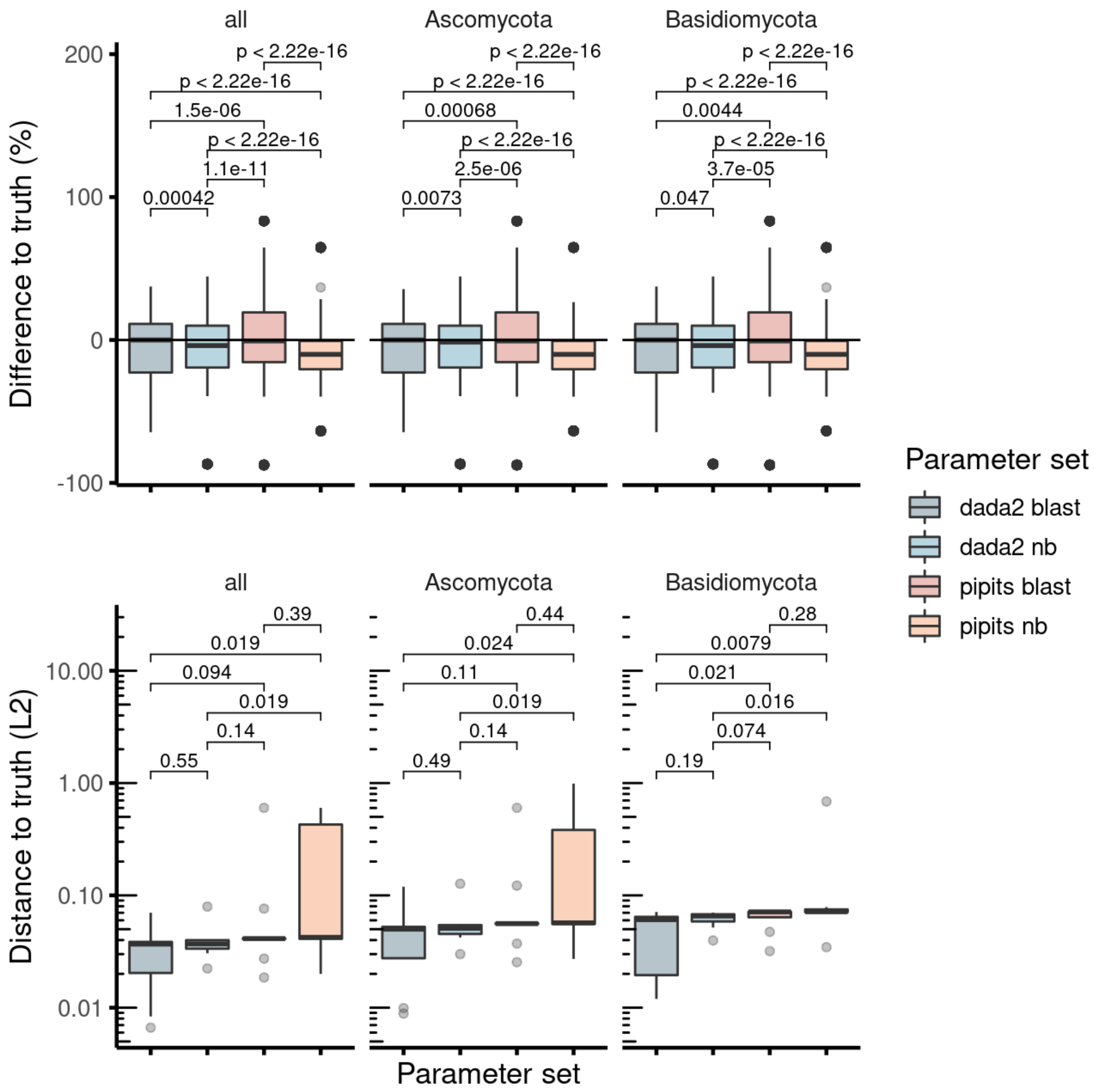


**Fig. S2:** Benchmarking taxon abundance. DAnIEL was run on 10 samples consisting simulated reads using denoising methods DADA2 and PIPITS and classification methods BLAST consensus (blast) and Naïve Bayes (nb). Difference of measured to true abundance was calculated for each sample and taxon. Furthermore, L2 norm of measured abundance profile to the true one was calculated for each sample. DADA2 in combination with BLAST yielded the most accurate results.
